# A somatosensory computation that unifies limbs and tools

**DOI:** 10.1101/2021.12.09.472009

**Authors:** Luke E. Miller, Cécile Fabio, Frédérique de Vignemont, Alice Roy, W. Pieter Medendorp, Alessandro Farnè

## Abstract

It is often claimed that tools are embodied by the user, but whether the brain actually repurposes its body-based computations to perform similar tasks with tools is not known. A fundamental computation for localizing touch on the body is *trilateration*. Here, the location of touch on a limb is computed by integrating estimates of the distance between sensory input and its boundaries (e.g., elbow and wrist of the forearm). As evidence of this computational mechanism, tactile localization on a limb is most precise near its boundaries and lowest in the middle. We show that the brain repurposes trilateration to localize touch on a tool. In a large sample of participants, we found that localizing touch on a tool produced the signature of trilateration, with highest precision close to the base and tip of the tool. A computational model of trilateration provided a good fit to the observed localization behavior. To further demonstrate the computational plausibility of repurposing trilateration, we implemented it in a three-layer neural network that was based on principles of probabilistic population coding. This network determined hit location in tool-centered coordinates by using a tool’s unique pattern of vibrations when contacting an object. Simulations demonstrated the expected signature of trilateration, in line with the behavioral patterns. Our results have important implications for how trilateration may be implemented by somatosensory neural populations. We conclude that trilateration is a fundamental spatial computation that unifies limbs and tools.

## Introduction

The proposal that the brain treats a tool as if it were an extended limb (tool embodiment) was first made over a century ago (Head and Holmes, 1911). From the point of view of modern neuroscience, embodiment would entail that the brain reuses its sensorimotor computations when performing the same task with a tool as with a limb. There is indirect evidence that this is the case (for reviews, see Maravita and Iriki, 2004; Martel et al., 2016), such as the ability of tool-users to accurately localize where a tool has been touched (Miller et al., 2018) just as they would on their own body. Several studies have highlighted important similarities between tool-based and body-based tactile spatial processing (Yamamoto and Kitazawa, 2001; Kilteni and Ehrsson, 2017; Miller et al., 2018), including at the neural level in the activity of frontoparietal regions (Miller et al., 2019; Pazen et al., 2020; Fabio et al., 2021). Tool use also modulates somatosensory perception and action processes (Cardinali et al., 2009; Cardinali et al., 2011; Cardinali et al., 2012; Sposito et al., 2012; Canzoneri et al., 2013; Miller et al., 2014; Garbarini et al., 2015; Cardinali et al., 2016; Miller et al., 2017; Martel et al., 2019; Romano et al., 2019; Miller et al., 2019b). While these findings suggest functional similarities between tools and limbs, direct evidence that body-based computational mechanisms are repurposed to sense and act with tools is lacking. The present study uses tool-based sensing as a case study to provide the first neurocomputational test of embodiment.

Tactile localization on the body is often characterized by greater precision near body-part boundaries (e.g., joints or borders), a phenomenon called perceptual anchoring (Cholewiak and Collins, 2003; de Vignemont et al., 2009). We recently found converging evidence that perceptual anchors are the signature of *trilateration* (Miller et al., 2022), a computation used by surveyors to localize an object within a map. To do so, a surveyor estimates the object’s distance from multiple landmarks of known positions. When applied to body maps (Figure 1A, bottom), a ‘neural surveyor’ localizes touch on a body part by estimating the distance between sensory input and body-part boundaries (e.g., the wrist and elbow for the forearm). To estimate the touch location in limb-centered coordinates, these two distance estimates can be integrated to produce a Bayes-optimal location percept (Ernst and Banks, 2002; Kording and Wolpert, 2004; Clemens et al., 2011). Consistent with Weber’s Law (Petzschner et al., 2015), we found that the noise in each distance estimate increased linearly as a function of distance (Figure 1B). Integrating them resulted in an inverted U-shaped noise profile across the surface, with the lowest noise near the boundaries and highest noise in the middle (i.e., perceptual anchoring).

**Figure 1.**
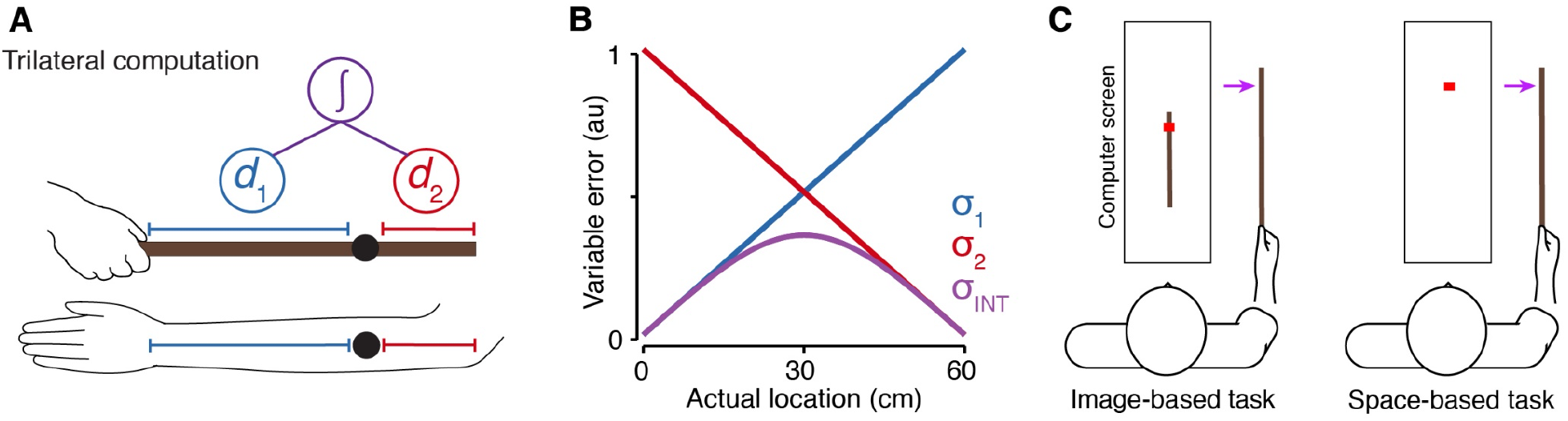
Model of trilateration and tool-sensing paradigm. (A) The trilateral computation applied to the space of the arm (bottom) a hand-held rod (top). Distance estimates from sensory input (black) and each boundary (D1 and D2) are integrated (purple) to form a location estimate. (B) In our model, the noise in each distance estimate (D1, D2) increases linearly with distance. The integrated estimate forms an inverted U-shaped pattern. (C) Two tool-sensing tasks used to characterize tactile localization on a hand-held rod. The purple arrow corresponds to the location of touch in tool-centered space. The red square corresponds to the judgment of location within the computer screen.

In the present study, we investigated whether trilateration is repurposed to localize touch a tool (Figure 1A). If this is indeed the case, localizing touch on a tool would be characterized by an inverted U-shaped pattern of variable errors across its surface (Figure 1B). We first provide a theoretical formulation of trilateration, arguing that the brain uses the tool’s vibrational properties to stand-in for a representation for the physical space of a tool (Miller et al., 2018); the brain could therefore repurpose trilateration by computing over a vibratory feature space (Figure 2). In this formulation (see Methods for more details), its boundaries stand in for the boundaries of tool-centered space and distance estimates (Figure 1A) are computed within a neural representation of the feature space. We then characterize the ability of participants to localize touch on a tool (Figure 1C) and use computational modelling to verify the expected computational signature of trilateration. Finally, we use neural network modelling to implement the vibration-to-location transformation, further solidifying the plausibility of trilaterating touch location on a tool.

**Figure 2.**
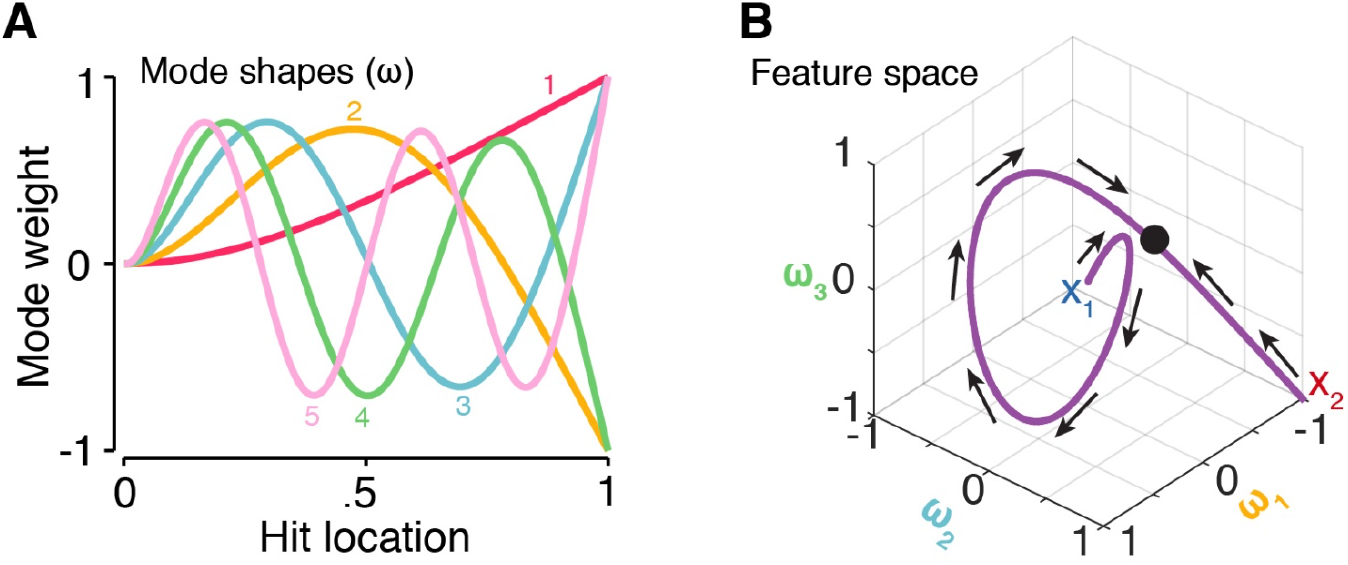
Vibration modes and feature space. (A) The shape of the first five modes *ω* for contact on a cantilever rod. The weight of each mode varies as a function of hit location. Each hit location is characterized by a unique combination of mode weights. (B) The vibration-location feature space (purple) from handle (X_1_) to tip (X_2_). This feature space is isomorphic with the actual physical space of the rod. *ω* corresponds to a resonant frequency, the black dot corresponds to the hit location (as in Figure 1A) within the feature space, and the arrows are the gradients of distance estimation during trilateration.

## Material and Methods

### Theoretical formulation of trilateration

In the present section, we provide a theoretical formulation of trilateration and how it can be applied to localizing touch within a somatosensory-derived coordinate system, be it centered on a body part or the surface of a tool (Figure 1A). The general *computational goal* of trilateration is to estimate the location of an object by calculating its distance from vantage points of known position, which we will refer to as landmarks. Applied to tactile localization, this amounts to estimating the location of touch by averaging over distance estimates taken from the boundaries of the sensory surface (Figure 1A), which serve as the landmarks and are assumed to be known to the nervous system via either learning or sensory feedback (Longo et al., 2010). For a body part (e.g., forearm), the landmarks are often its joints (e.g., wrist and elbow) and lateral sides. For simple tools such as rods, the landmarks correspond to their handle and tip—previous research has shown that users can sense their positions from somatosensory feedback during wielding (Debats et al., 2012).

We will first consider the general case of localizing touch within an unspecified somatosensory coordinate system. For simplicity, we will consider only a single dimension of the coordinate system, with localization between its two boundaries. We propose that the somatosensory system only needs three spatial variables, {*x*_1_, *x*_2,_ *x*_3_}, to derive an estimate 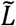 of the actual location of touch *L* in surface-centered coordinates. The variables *x*_1_ and *x*_2_ correspond to the proximal and distal boundaries, respectively. The variable *x*_3_ corresponds to the sensory input. Due to noise (Faisal et al., 2008), the nervous system does not represent variables as point estimates but as probability densities over some range of values (Pouget et al., 2013). Assuming normally-distributed noise, each variable *x*_*i*_ can be thus thought of as a Gaussian likelihood

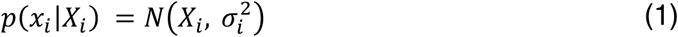

where the mean *X*_*i*_ corresponds to its true spatial position and the variance 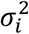 corresponds to the uncertainty in its internal estimate. Here, *X*_1_ and *X*_2_ are the true positions of the landmarks (i.e., boundaries) and *X*_3_ is the position of the sensory input. It is important to note here that these positions can be specified within any shared coordinate system. For example, touch on the body is thought to initially be represented in skin-based coordinates (Medina and Coslett, 2010), not coordinates centered on a limb. The relationship between *X*_3_ and *L* therefore remains ambiguous without the proper computation to transform it into the actual surface-centered coordinates (Longo et al., 2010).

Trilateration performs the necessary computation to transform *x*_3_ into surface-centered coordinates (Miller et al., 2022). It does so by calculating its distance from the proximal and distal boundaries of the coordinate system (*x*_1_ and *x*_2_, respectively), producing two additional estimates:

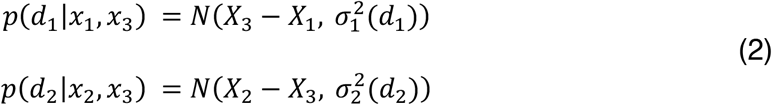

where each distance estimate *d*_*i*_ corresponds to a Gaussian likelihood with a mean equal to the distance between *X*_3_ and the respective boundary and a variance that scales with distance. That is, localization estimates are more precise when the touch is physically closer to a boundary than when it is farther away (Figure 1B). This distance-dependent noise is consistent with the Weber-Fechner law (Petzschner et al., 2015) and is a consequence of how distance computation is implemented by a neural decoder (see below).

Given the above distance estimates (Eq. 2), we can derive two estimates of touch location 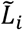 that are aligned within a common coordinate system:

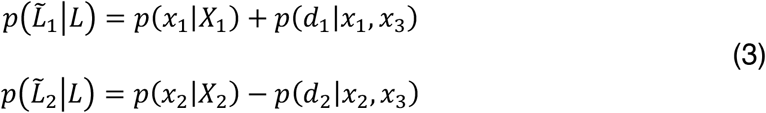

These two location estimates can be used to derive a final estimate. However, given the presence of distance-dependent noise, the precision of each estimate will vary across the sensory surface (Figure 1B). Assuming a flat prior for touch location, the statistically optimal solution (i.e., maximum likelihood) is to integrate both estimates:

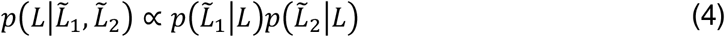

Here, the mean (*μ*_*INT*_) and variance 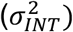 of the integrated surface-centered posterior distribution depend on the means (*μ*_1_ and *μ*_2_) variances (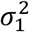 and 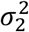) of the individual estimates:

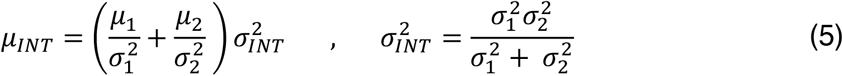

The integrated posterior 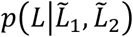 thus reflects the maximum-likelihood estimate of touch location *L*. Given that the noise in each individual estimate scales linearly with distance, integration has the consequence of producing an inverted U-shaped pattern of variance (Figure 1B). This pattern of variability serves as a *computational signature* of trilateration, which we have observed for tactile localization on the arm and fingers (Miller et al., 2022). The present study investigates whether this is the case for localizing touch on a hand-held rod. Our computational analyses implement this probabilistic model of trilateration (see below).

### Computing a tool-centered spatial code with trilateration

Let us now consider the more specific case of performing trilateration for touch on a tool (Figure 1A, top). Because the tool surface is not innervated, spatial information does not arise from a distribution of receptors but must instead be inferred from sensory information during tool-object contact. However, as we will see, this information forms a feature space that can computationally stand in for the real physical space of the tool (Figure 2). Trilateration can be performed on this feature space, leading to a tool-centered code.

As with the body, the somatosensory system needs three variables, {*x*_1_, *x*_2,_ *x*_3_}, to derive an estimate 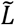 of the actual location of touch *L* in tool-centered coordinates. The representational nature of these variables depends on the type of sensory information that encodes where a tool was touched. We have previously argued that touch location is encoded in rod’s resonant frequencies (Miller et al., 2018). The frequencies of these modes are determined by physical properties of the rod, such as its length and material. However, the relative amplitudes of each mode is determined by touch location (Figure 2A), a pattern that is invariant across rods. The link between location and amplitude is captured by the shape of the modes.

Touch location can therefore be encoded in a unique combination of modal amplitudes, called *vibratory motifs*. These motifs form a multidimensional feature space that forms a vibration-to-location isomorphism (Figure 2B). Theoretically, this isomorphic mapping between the feature space of the vibrations and tool-centered space can computationally stand in for the physical space of the rod. We can therefore re-conceptualize the three initial spatial variables, {*x*_1_, *x*_2,_ *x*_3_}, in relation to the isomorphism. The estimates *x*_1_ and *x*_2_ encode the location of the proximal and distal boundaries within the feature space, respectively. The estimate *x*_3_ encodes the sensory input, which in our case is the vibration amplitude in each mode. Once the nervous system has learned the isomorphic mapping, the trilateral computation (Equations 2-5) can be used to derive an estimate of the tool-centered location of touch (Figure 2B). To concretely demonstrate this possibility, we implemented this isomorphic mapping in a simple neural network.

### Neural network implementation for trilateration on a tool

Somatosensory regions are characterized by spatial maps of the surface of individual body parts (Penfield and Boldrey, 1937). Based on this notion, we previously applied the above formulation of trilateration to tactile localization on the body surface, and implemented it in a biologically inspired two-layer feedforward neural network (Miller et al., 2022). The first layer consisted of units that were broadly tuned to touch location in skin-based coordinates, as is thought to be encoded by primary somatosensory cortex. The second layer consisted of units whose tuning was characterized distance-dependent gradients (either in peak firing rate and/or tuning width) that were anchored to one of the joints. They therefore embodied the distance computation as specified in Equations 2–3. A Bayesian decoder demonstrated that the behavior of this network matched what would be expected by optimal trilateration (Equations 2–5), displaying distance-dependent noise and an inverted U-shaped variability following integration.

While this network relies on the observation that individual primary somatosensory neurons are typically tuned to individual regions of the skin (Delhaye et al., 2018), can it also be re-used for performing trilateration in vibration space? The vibratory motifs are unlikely to be spatially organized across the cortical surface. Instead, the nervous system must internalize the isomorphic mapping between the motifs and the physical space of the tool (Figure 2). We have previously found that disrupting the expected vibrations disrupts localization (Miller et al., 2018), suggesting that the user has internal models of rod dynamics (Imamizu et al., 2000). We assume that this internal model is implemented in units that are tuned to the statistics of the vibratory motifs.

We implemented the trilateral computation (Equations 2–5) in a three-layer neural network with four processing stages (Figure 3): First, the amplitudes of each mode are extracted by a population of units with subpopulations tuned to each resonant mode (Layer 1). Second, activation in each subpopulation is integrated by units tuned to the multidimensional statistics of the motifs (Layer 2). This layer effectively forms the internal model of the feature space that is isomorphic to the rod’s physical space. Next, this activation pattern is transformed into tool-centered coordinates (Equations. 2–3) via two decoding subpopulations whose units are tuned to distance from the boundaries of the feature space (Eq. 3; Layer 3). The population activity of each decoding subpopulations reflects the likelihoods in Equation 4 (Jazayeri and Movshon, 2006). Lastly, the final tool-centered location estimate is derived by a Bayesian decoder (Ma et al., 2006) that integrates the activity of both subpopulations (Eq. 5).

**Figure 3.**
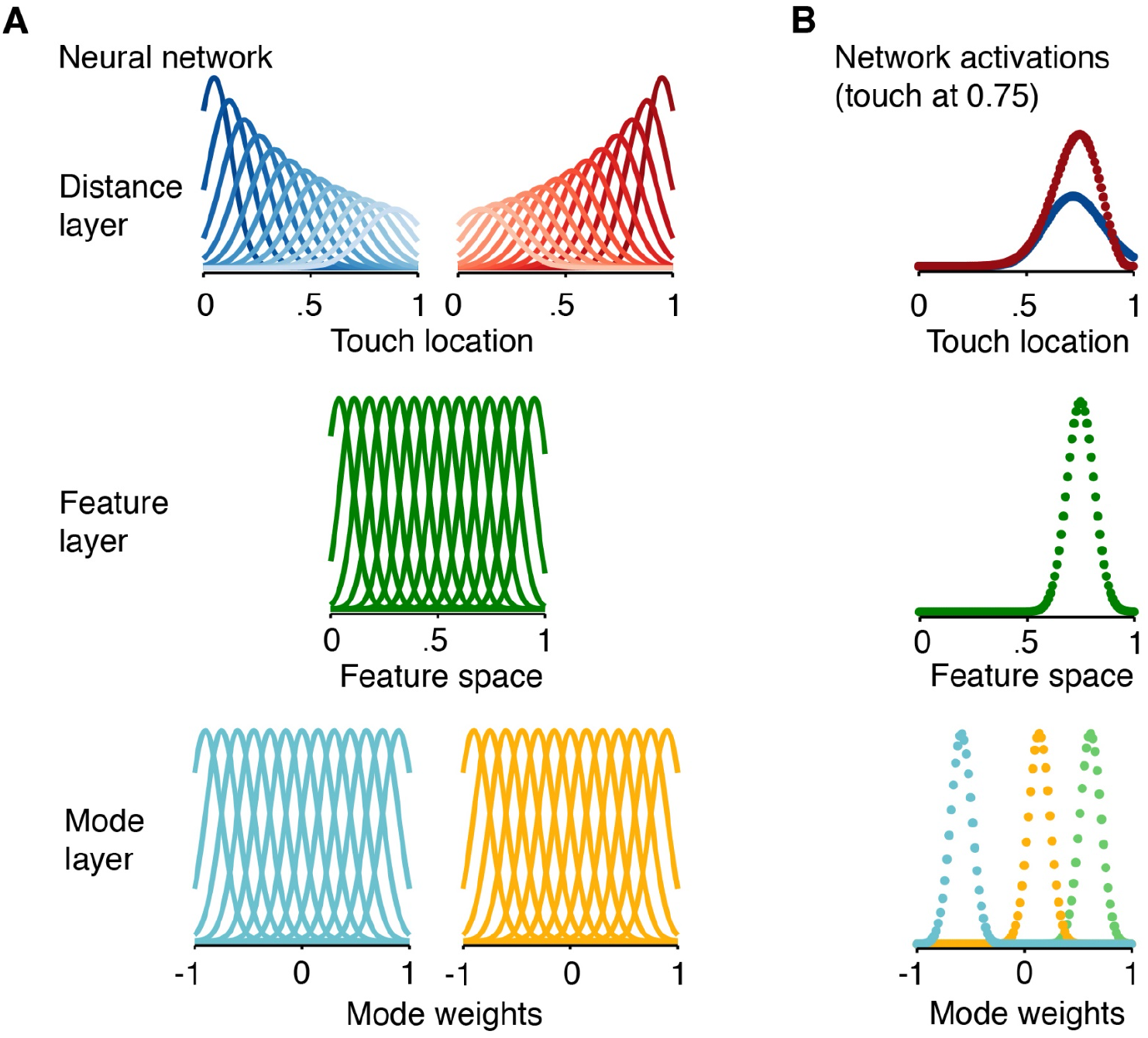
Neural network implementation of trilateration. (A) Neural network implementation of trilateration: (lower panel) the Mode layer is composed of sub-populations (two shown here) sensitive to the weight of individual modes (see Figure 2A), which are location-dependent; (middle panel) the Feature layer takes input from the model layer and encodes the feature space (see Figure 2B), which forms the isomorphism with the physical space of the tool; (upper panel) the Distance layer is composed of two subpopulations of neurons with distance-dependent gradients in tuning properties (shown: firing rate and tuning width). The distance of a tuning curve from its “anchor” is coded by the luminance, with darker colors corresponding to neurons that are closer to the spatial boundary. (B) Activations for each layer of the network averaged over 5000 simulations when touch was at 0.75 (space between 0 and 1). Each dot corresponds to a unit of the neural network. (lower panel) mode layer, with three of five subpopulations shown; (middle panel) feature layer; (upper panel) distance layer of localization for each decoding subpopulation.

The feature space of vibrations is multidimensional, being composed of a theoretically infinite number of modes. However, only the first five modes (Figure 2A) are typically within the bandwidth of mechanoreceptors (i.e., ∼10-1000 Hz; Johansson and Flanagan, 2009). The first layer of our network was therefore composed of units tuned to the amplitudes of the modes (Figure 3A, bottom). This layer was composed of five subpopulations, one for each mode. These units were broadly-tuned with Gaussian (bell-shaped) tuning curves *f*^*M*^ of the following form:

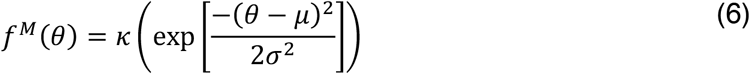

where *κ* is the peak firing rate (i.e., gain), *μ* is the tuning center related to the amplitude of the specific mode, *θ* is the mode amplitude of the stimulus, and *σ*^2^ is the variance of the tuning curve. We modelled the response properties of these units for a given contact location on the rod with likelihood functions 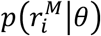 denoting the probability that mode amplitude *θ* caused 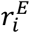 spikes in encoding unit *i*. The likelihood function 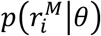 was modeled as a Poisson probability distribution with a Fano factor of one according to the following equation:

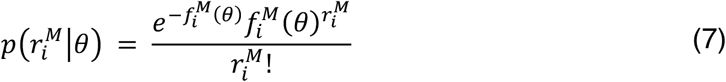

where 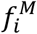 is the tuning curve of unit *i*. The population response of the encoding units is denoted by a vector 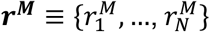, where 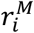 is the spike count of unit *i*.

The amplitude *θ* of each mode is tied directly to the stimulus location *L* (Miller et al., 2018). The function of the next layer is to integrate the amplitudes of each mode, encoded in ***r***^***M***^, into a representation of the feature space that can be directly linked to *L*. It does so via units with bell-shaped tuning curves *f*^*S*^ over the feature space (Figure 3A, middle). The population activity ***r***^***S***^ of this layer is a combination of (1) the synaptic input *W*^*S*^ · ***r***^***M***^, where ‘·’ is the dot product and *W*^*S*^ is the matrix of all synaptic weights; and (2) the uninherited Poisson noise in the decoding unit’s spiking behavior (Eq. 7). Each unit *i* in the second layer was fully connected to each unit in the first layer via a vector synaptic weights 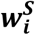, which was set to be proportional to ***r***^***M***^ for each touch location *L*. For simplicity, the input into the second layer (***f***^***S***^(*j*)) corresponded to the winner-take-all of the synaptic input 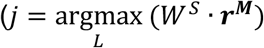.

The function of the third layer was to estimate the location of *L* in tool-centered coordinates given the population response ***r***^***S***^ in the feature space layer. We implemented this computation in two independent decoding subpopulations, each of which was “anchored” to one of the boundaries of the feature space (Figure 3A, top). The population activity ***r***^***D***^ of each subpopulation corresponded to: 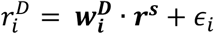, where 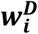 is the vector of synaptic weights connecting unit *i* to the second layer and *ϵ*_*i*_ is the uninherited Poisson noise in the unit’s spiking behavior (Eq. 7). Each unit in the decoding layer was fully connected to each unit in the encoding layer via ***w***^***D***^. We used the Matlab function *fmincon* to find the positive-valued weight vector that produced the decoding unit’s pre-specified tuning curve (see below).

As in our previous neural network for body-centered tactile localization (Miller et al., 2022), the distance computation (Equations 2–3) was embodied by distance-dependent gradients in the tuning of units *f*^*D*^ in each decoding subpopulation. The gain *κ* of these units formed a distance-dependent gradient (close-to-far: high-to-low gain) across the length of the feature space.

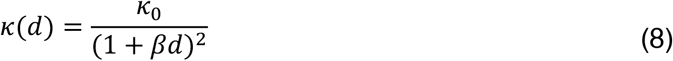

where *κ*_0_ corresponds to the gain of the tuning curve centered on the landmark’s location (i.e., distance zero), *d* is the distance from the center of the tuning curve (*d* ≥ 0) and the landmark, and *β* is a scaling factor. The width *σ* of each tuning curve can be uniform in either linear or log space. In the latter case, tuning width also forms a distance-dependent gradient (close-to-far: narrow-to-wide tuning) in linear space (Nieder and Miller, 2003), consistent with the Weber-Fechner law.

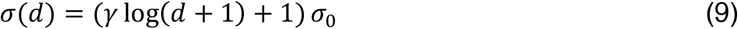

where *σ*_0_ corresponds to the width of the tuning curve centered on the landmark’s location, *d* is the distance from the center of the tuning curve and the landmark (*d* ≥ 0), and *γ* is a scaling factor. It is important to note that these units *f*^*D*^ are tuned to the feature space, not the vibrations themselves (as in the encoding layer). Given the isomorphism, we can therefore link their response properties directly to the location of touch *L*.

When neuronal noise is Poisson-like (as in Eq. 7), the gain of a neural population response reflects the precision (i.e., inverse variance) of its estimate (Ma et al., 2006). Therefore, given the aforementioned distance-dependent gradient in gain, noise in each subpopulation’s location estimate (that is, its uncertainty) will increase as a function of distance from a landmark (i.e., the handle or tip). Consistent with several studies (Jazayeri and Movshon, 2006; Ma et al., 2006), we assume that the population responses encode log probabilities. We can therefore decode a maximum likelihood estimates of each subpopulation as follows:

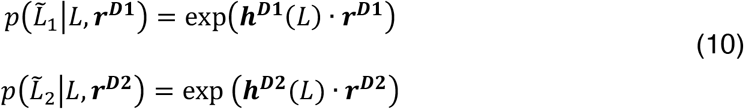

where ***h***^***D***^ is a kernel and ***r***^***D***^is the subpopulation response. When neural responses are characterized by independent Poisson noise (Eq. 7), ***h***^***D***^ is equivalent to the log of each subpopulation’s tuning curve ***f***^***D***^ at value *L* (Jazayeri and Movshon, 2006; Ma et al., 2006). Assuming that the population response reflects log probabilities, optimally integrating both estimates (Eq. 5) amounts to simply summing the activity of each subpopulation.

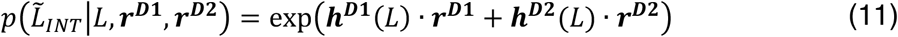

where the optimal estimate 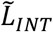 on a given trial *n* can be written as the location for which the log-likelihood of the summed population responses is maximal.

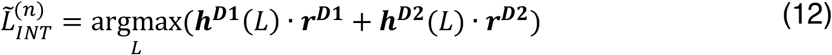

We previously demonstrated that the above neural network, with a different encoding layer, implements trilateration for localizing touch in body-centered coordinates. Our present neural network (Equations 6–12) generalizes the Bayesian formulation of trilateration (Equations 2–5) to localizing touch on a tool, using a vibratory feature space as a proxy for tool-centered space. The flow of activity in this network can be visualized at Figure 3B, where the touch occurs at 75% the surface of the tool. To systematically investigate the behavior of this network, we simulated 5000 instances of touch at wide range of locations (10% to 90% of the space) on the tool surface using the above network. The values for the above parameters in all layers can be seen in Table 1. All units of each layer shared the same parameter values. We used a maximum log-likelihood decoder to localize touch from the overall response of each subpopulation separately or integrated.

**Table 1.** Initial neural network parameter values.

| | $f^M$ | $f^S$ | $f^{D1}$ | $f^{D2}$ |
| --- | --- | --- | --- | --- |
| $\mu$ | -1.5:0.02:1.5 | -40:1:140 | 0:1:140 | -40:1:100 |
| $\kappa$ or $\kappa_0$ | 25 | 25 | 25 | 25 |
| $\sigma$ or $\sigma_0$ | 0.08 | 3.40 | 3.40 | 3.40 |
| $\beta$ | — | — | 0.01 | 0.01 |
| $\gamma$ | — | — | 0.5 | 0.5 |

### Behavioral Experiment

#### Participants

Forty right-handed participants (24 females, 23.65 ± 2.48 years of age) in total completed our behavioral experiments. Two participants were removed due to inability to follow task instructions, leaving thirty-eight in total to be analyzed. All participants had normal or corrected-to-normal vision and no history of neurological impairment. Every participant gave informed consent before the experiment. The study was approved by the ethics committee (CPP SUD EST IV, Lyon, France).

#### Experimental procedure

During the task, participants were seated comfortably in a cushioned chair with their torso aligned with the edge of a table and their right elbow placed in a padded arm rest. The entire arm was hidden from view with a long occluding board. A 60 cm-long rod (handle length: 12-cm; cross-sectional radius: 0.75 cm) was placed in their right hand. This rod was either wooden (twenty-five participants) or PVC (thirteen participants). The arm was placed at a height necessary for a 1 cm separation between the object (see below) and the rod at a posture parallel with the table. On the surface of the table, an LCD screen (70 × 30 cm) lay backside down in the length-wise orientation; the edge of the LCD screen was 5 cm from the table’s edge. The center of the screen was aligned with the participant’s midline.

The task of participants was to localize touches resulting from active contact between the rod and an object (foam-padded wooden block). In an experimental session, participants completed two tasks with distinct reporting methods (order counterbalanced across participants). In the *image-based task*, participants used a cursor to indicate the corresponding location of touch on a downsized drawing of a rod (20 cm in length; handle to tip); the purpose of using a downsized drawing was to dissociate it from the external space occupied by the real rod. The drawing began 15 cm from the edge of the table, was raised 5 cm above the table surface, and was oriented in parallel with the real rod. The red cursor (circle, 0.2 cm radius) was constrained to move in the center of the screen occupied by the drawing. In the *space-based task*, participants used a cursor to indicate the corresponding location of touch within in an empty LCD screen (white background). The cursor was constrained to move along the vertical bisection of the screen and could be moved across the entire length of the screen. It is critical to note that in this task, participants were forced to rely on somatosensory information about tool length and position as no other sensory cues were available to do so.

The trial structure for each task was as follows: In the ‘Pre-contact phase’, participants sat facing the computer screen with their left hand on a trackball. A red cursor was placed at a random location within the vertical bisection of the screen. A ‘go’ cue (brief tap on the right shoulder) indicated that they should actively strike the object with the rod. In the ‘Localization phase’, participants made their task-relevant judgment with the cursor, controlled by the track-ball. Participants never received feedback about their performance. To minimize auditory cues during the task, pink noise was played continuously over noise-cancelling headphones.

The object was placed at one of six locations, ranging from 10 cm from the handle to the tip (10–60 cm from the hand; steps of 10 cm). The number of object locations was unknown to participants. In each task, there were ten trials per touch location, making 60 trials per task and 120 trials in total. The specific location for each trial was chosen pseudo-randomly. The entire experimental session took approximately 45 minutes.

The experiment started with a five-minute sensorimotor familiarization session. Participants were told to explore, at their own pace, how the tool felt to contact the object at different locations. They were instructed to pay attention to how the vibrations varied with impact location. Visual and auditory feedback of the tool and tool-object contact was prevented with a blindfold and pink noise, respectively. Participants were, however, allowed to hold the object in place with their left hand while contacting it with the tool but were not allowed to haptically explore the rod.

At the end of the space-based task, participants used the cursor to report where they felt the tip of the rod (aligned in-parallel to the screen). The judged location of the tip (mean: 56.5 cm; SEM: 1.62 cm) was very similar to the rod’s actual length (i.e., 60 cm). It is critical to reiterate here that participants had never seen the rod prior up to this point of the experiment, and likely relied on somatosensory feedback about its dimensions.

### Data Analysis

#### Regression analysis

Prior to analysis, all judgments in the image-based task were converted from pixels of drawing space to percentage of tool space. All judgments in the space-based task were normalized such that their estimated tip location corresponded to 100% of tool space. We then used least-squares linear regression to analyze the localization accuracy. The mean localization judgment for each touch location was modelled as a function of actual object location. Accuracy was assessed by comparing the group-level confidence intervals around the slope and intercept.

#### Trilateration model

Our model of trilateration in the somatosensory system assumes that the perceived location of touch is a consequence of the optimal integration of two independent location estimates, 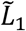 and 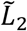. This is exemplified in our formulation of trilateration (Equations 1-5). Trilateration predicts that noise in each estimate varies linearly as a function of the distance of touch from two landmarks (Equation 2; Figure 1B), corresponding to the handle and tip. For any location of touch *L* along a tactile surface, the variance in each landmark-specific location estimate 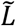 can therefore be written as follows:

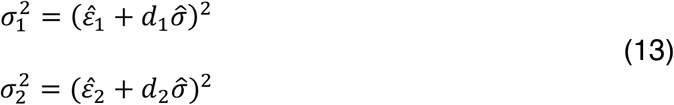

in which 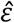 is a landmark-specific intercept term that likely corresponds to uncertainty in the location of each landmark, *d* is the distance of touch location *L* from the landmark (Equations 2–3), and 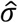 is the magnitude of noise per unit of distance. We assume that the noise term 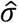 corresponds to a general property of the underlying neural network and therefore model it as the same value for each landmark. The distance-dependent noise for the integrated estimate is therefore:

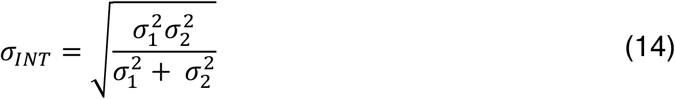

The three parameters in the model (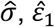, and 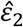) are properties of the underlying neural processes that implement trilateration and are therefore not directly observable. They must therefore be inferred using a reverse engineering approach, where they serve as free parameters that are fit to each participant’s variable errors. We simultaneously fit the three free parameters to the data using non-linear least squares regression. Optimal parameter values were obtained through maximum likelihood estimation using the Matlab routine *fmincon*. All modelling was done with the combined data from both localization tasks. *R*^*2*^ values for each participant in each experiment were taken as a measure of the goodness-of-fit between the observed and predicted pattern of location-dependent noise.

#### Boundary truncation model

Boundary truncation provides one alternative model to trilateration This model assumes that the estimate of location 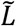 corresponds to a Gaussian likelihood whose variance is *identical* at all points on the rod. The inverted U-shaped variability arises because these likelihoods are truncated by a boundary, either by the range of possible responses or by a categorical boundary (e.g., between handle and tip). As in Equation 1, we can model each likelihood 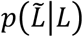 as a normal distribution *N*(*μ*_*L*_, σ_*L*_) where *μ*_*L*_ is the location of touch *L* and σ_*L*_ is the standard deviation. The posterior estimate 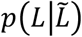 then corresponds to a likelihood truncated at *γ*_1_ and *γ*_2_, where *γ*_2_ > *γ*_1_. Doing so will distort the mean and variance of the posterior estimate.

We fit this truncation model to the participant-level variable errors in each of our experiments. The standard deviation for each location, σ_*T*_(*L*), was determined by truncating a normal distribution at *γ*_1_ and *γ*_2_ using the *makedist* and *truncate* functions in MATLAB. The model therefore had three free parameters, σ_*T*_, *γ*_1_ and *γ*_2_. The value of σ_*T*_ was constrained between 1 and 40; *γ*_1_ between −30 and 30; and *γ*_2_ between 70 and 130 (units: % of rod surface). These ranges—particularly for *γ*_1_ and *γ*_2_—are quite unrealistic, but were chosen to maximize a good fit with the variable errors. Fitting procedures for this model were the same as the trilateration model.

#### Model comparisons

We used the Bayesian Information Criterion (BIC) to compare the boundary and trilateration models. The difference in the BIC (ΔBIC) was used to determine a significant difference in fit. Consistent with convention, the chosen cutoff for moderate evidence was a ΔBIC of 2 and the cutoff for strong evidence was a ΔBIC of 6.

## Results

### Accurate localization of touch on a tool

In the current experiment (n=38), we investigated whether tactile localization on a 60-cm hand-held rod is characterized by the U-shaped pattern of variability (Figure 1B) that is characteristic of trilateration when localizing touch on the body. In two tasks, we measured participants’ ability to localize an object that was actively contacted with a hand-held tool. In the *image-based task*, participants indicated the point of touch on a downsized drawing of the tool. In the *space-based task*, participants indicated the point of touch in external space. The latter task ensured that localization was not truncated by boundaries in the range of possible responses.

Consistent with prior results (Miller et al., 2018), we found that participants were generally quite accurate at localizing touch on the tool. Linear regressions (Figure 4A) comparing perceived and actual hit location found slopes near unity both the image-based task (mean slope: 0.93, 95% CI [0.88, 0.99]) and the space-based task (mean slope: 0.89, 95% CI [0.82, 0.95]). Analysis of the variable errors (2×6 repeated measures ANOVA) found a significant main effect of hit location (*F*(5,185)=36.1, *p*<.001) but no main effect of task (*F*(1,37)=0.39, *p*=.54) or an interaction (*F*(5,185)=0.21, *p*=.96). Crucially, the pattern of variable errors (Figure 4B) in both tasks displayed the hypothesized inverted U-shape, which was of similar magnitude to what we observed for touch on the arm (Cholewiak and Collins, 2003; Miller et al., 2022).

**Figure 4.**
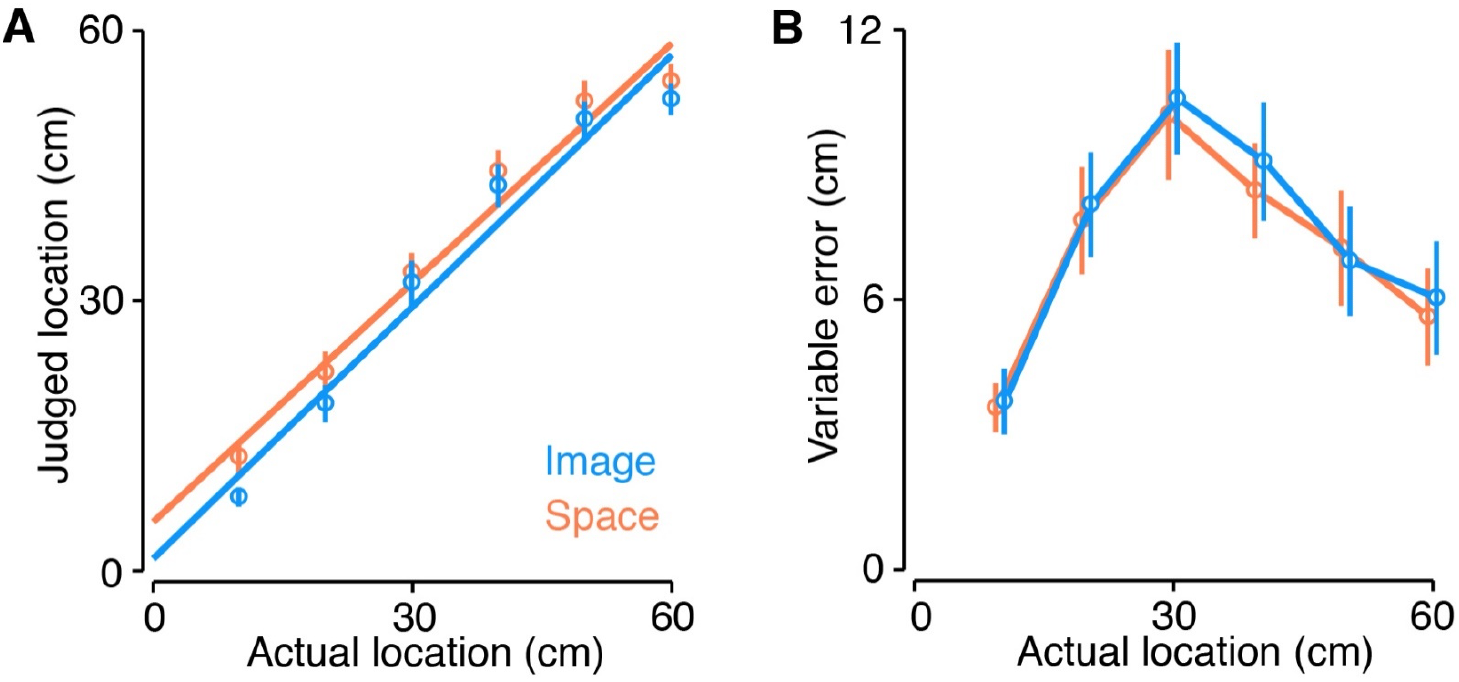
Localization and variable error for both tasks. (A) Regressions fit to the localization judgments for both the image-based (blue) and space-based (orange) tasks. Error bars correspond to the group-level 95% confidence interval. (B) Group-level variable errors for both tasks. Error bars correspond to the group-level 95% confidence interval.

### Computational modelling of behavior

We next used computational modelling to confirm that the observed pattern of variable errors was indeed due to trilateration. We fit each participant’s variable errors with a probabilistic model of optimal trilateration (Figure 1A-B) that was derived from its theoretical formulation (see Methods). We compared the trilateration model to an alternative hypothesis: The inverted U-shaped pattern is due to truncation at the boundaries of localization (Petzschner et al., 2015), which cuts off the range of possible responses and thus produces lower variability at these boundaries. We fit a boundary truncation model to directly compare to our trilateration model. Given the lack of a main effect of task and to increase statistical power, we collapsed across both tasks in this analysis.

Our computational model of trilateration provided a good fit to the variable errors observed during tactile localization on a tool. This was evident at the group-level, where the magnitude of variable errors was similar to what has been found when localizing touch on the arm (Figure 5A). We further observed a high coefficient of determination at the level of individual participants (mean *R*^*2*^: 0.71; range: 0.29–0.95); indeed, 30 out of 38 participants had an *R*^*2*^>0.6. The fits of the trilateration model to the data of 6 randomly chosen participants can be seen in Figure 5B. In contrast, the *R*^*2*^ of the boundary truncation model was substantially lower than the trilateration model (mean: 0.29; range: −0.19–0.71), never showing a better fit to the data in any participant.

**Figure 5.**
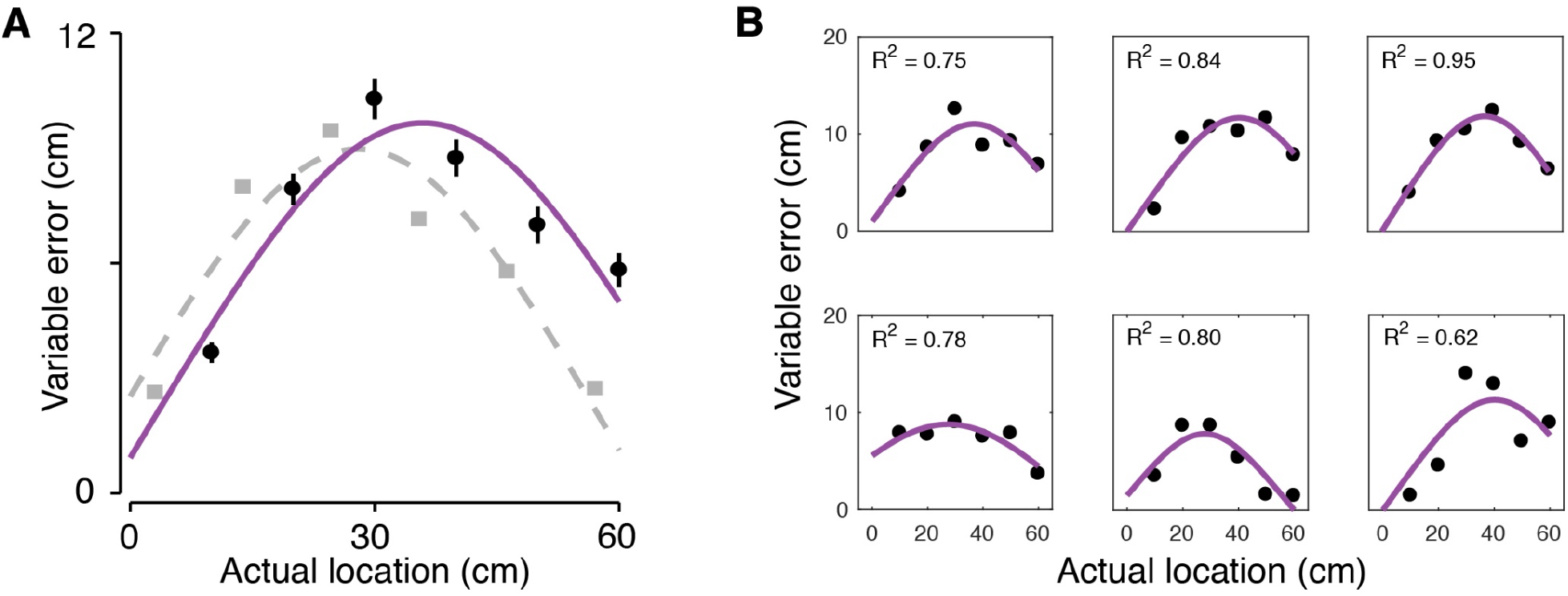
Trilateration model provides a good fit to localization behavior. (A) Fit of the trilateration model to the group-level variable error (black dots). The purple line corresponds to the model fit. The light gray line and squares correspond to variable errors for localization on the arm observed in Miller et al (2022); note that this data is size adjusted to account for differences in arm and rod size. (B) Fit of the trilateration model to the variable errors of six randomly chosen participants.

We next compared each model directly using the Bayesian Information Criteria (BIC). The BIC score for the trilateration model was lower in all 38 participants (mean±sd; Trilateration: 11.88±5.88; Truncation: 18.74±4.70). Statistically, 32 participants showed moderate evidence (ΔBIC>2) and 20 participants showed strong evidence (ΔBIC>6) in favor of trilateration. In total, our results strongly suggest that, as with the body, touch on a tool is localized via trilateration.

### Neural network simulations

Finally, we simulated trilateration on a tool using a biologically inspired neural network with a similar architecture as we have done previously. The goal of these simulations was to concretely demonstrate that the feature space of vibratory motifs could stand in for the physical space of the rod. Our neural network thus took the mode amplitudes as input and trilaterated the resulting touch location in tool-centered coordinates (5000 simulations per location).

Both subpopulations in the distance-computing layer (Layer 3; Figure 3, top) were able to localize touch with minimal constant error (Figure 6A, upper panel), demonstrating that each could produce unbiased estimates of location from the sensory input. However, as predicted given the gradient in tuning parameters, the noise in their estimates rapidly increased as a function of distance from each landmark (Figure 6B upper panel), forming an X-shaped pattern across the surface of the tool.

**Figure 6.**
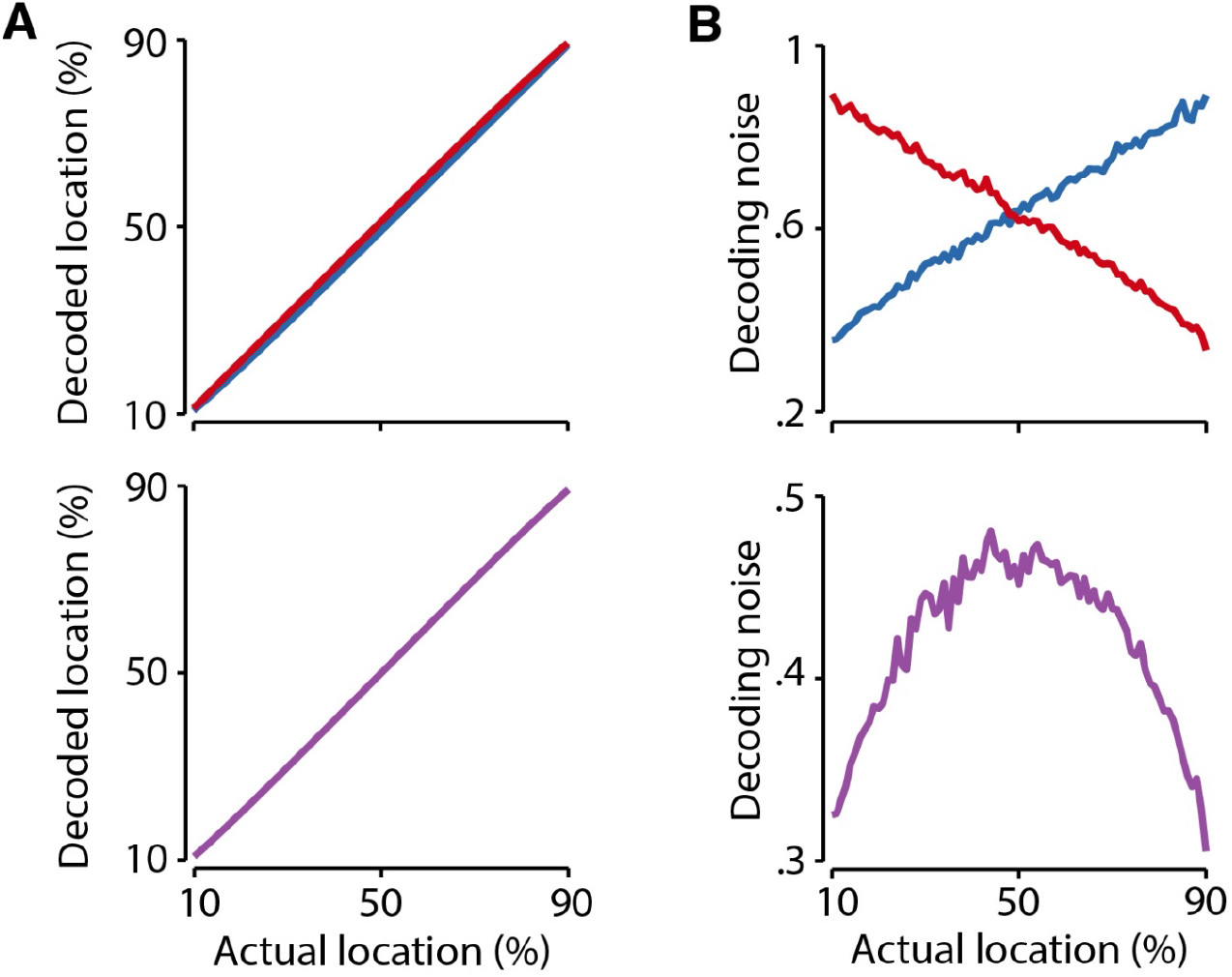
Neural network simulations. (A) Localization accuracy for the estimates of each decoding subpopulation (upper panel; L_1_, blue; L_2_, red) and after integration by the Bayesian decoder (lower panel; L_INT_, purple). (B) Decoding noise for each decoding subpopulation (upper panel) increased as a function of distance from each landmark. Note that distance estimates are made from the 10% and 90% locations for the first (blue) and second (red) decoding subpopulations, respectively. Integration via the Bayesian decoder (lower panel) led to an inverted U-shaped pattern across the surface. Note the differences in the y-axis range for both panels.

We next examined the output of the Bayesian decoder from Equations 11–12 (Figure 6, lower panel). As expected, we observed the computational signature of trilateration. Integrating both estimates resulted in an inverted U-shaped pattern of decoding noise across the surface of the tool (Figure 6B, lower panel), with the lowest decoding noise near the land-marks and the highest decoding variance in the middle. Crucially, this is the exact pattern of variability we observed in our behavioral experiments (see above) and have previously observed for tactile localization on the body. These simulations establish the plausibility of trilateration as a computation that can turn a vibratory code into a spatial representation.

## Discussion

If tools are embodied by the sensorimotor system, we would expect that the brain repurposes its body-based sensorimotor computations to perform similar tasks with tools. Using tactile localization as our case study, we uncovered multiple pieces of evidence that are consistent with this embodied view. First, as is the case for body parts, we observed that localizing touch on the surface of a tool is characterized by perceptual anchors at the handle and tip (de Vignemont et al., 2009). Second, computational modeling of behavioral responses suggests that they are the result of the probabilistic computation involving trilateration. Indeed, perceptual anchors are a computational signature of trilateration. Finally, using a simple three-layer population-based neural network, we demonstrated the possibility of trilateration in the vibratory feature space evoked by touches on tools. This neural network transformed a vibration-based input into a spatial code, reproducing perceptual anchors on the tool surface. These findings go well-beyond prior research on embodiment (Martel et al., 2016) by identifying a computation that functionally unifies tools and limbs. Indeed, they demonstrate that trilateration is the spatial computation employed by the somatosensory system to localize touch on body parts and tools alike (Miller et al., 2022). They further have important implications for how trilateration would be repurposed at a neural level for tool-extended sensing.

If trilateration is a fundamental spatial computation used by the somatosensory system it should be employed to solve the same problem (i.e., localization) regardless of whether the sensory surface is the body or a tool. Previous tactile localization studies have reported increased perceptual precision near the boundaries of the hands (Elithorn et al., 1953; Miller et al., 2022), arm (Cholewiak and Collins, 2003; de Vignemont et al., 2009; Miller et al., 2022), feet (Halnan and Wright, 1960), and abdomen (Cholewiak et al., 2004). These perceptual anchors are a signature of a trilateration computation (Miller et al., 2022). The results of the present study are consistent with the use of trilateration to localize touch on tools as well.

Our findings provide computational evidence that tools are *embodied* in the sensorimotor system (Martel et al., 2016), an idea that was proposed over a century ago (Head and Holmes, 1911). The close functional link between tools and limbs is not just a superficial resemblance but rather a reflection of the repurposing of neurocomputational resources dedicated to sensing and acting with a limb to that with a tool (Makin et al., 2017). This repurposing may be one reason that tool use leads to measurable changes in body perception and action processes (Canzoneri et al., 2013; Cardinali et al., 2009; Miller et al., 2014; Miller et al., 2019a).

There are, of course, important differences between limbs and tools. While the skin is innervated with sensory receptors, the somatosensory system must ‘tune into’ a tool’s mechanical response in order to extract meaningful information from it. We have previously proposed that where a rod is touched is encoded by the amplitudes of its resonant responses when contacting an object (Miller et al., 2018; Miller et al., 2019b). These resonant modes form a feature space that is isomorphic with the physical space of the tool. At a peripheral level, these resonances are re-encoded by the spiking patterns of tactile mechanoreceptors (Johnson, 2001). Therefore, unlike for touch on the body, localizing touch on a tool requires the somatosensory system to perform a temporal-to-spatial transformation.

We used neural network simulations to embody the necessary transformations to implement trilateration on a tool. Our neural network assumes that the human brain contains neural populations that encode for the full feature space of rod vibration. While very little is known about how these types of naturalistic vibrations are represented by the somatosensory system, our modeling results and prior research (Miller et al., 2018; Miller et al., 2019) suggest that there are neural populations that encode their properties. Previous work demonstrated that individual neurons in primary somatosensory cortex multiplex both amplitude and frequency in their firing properties (Harvey et al., 2013). Recent evidence further suggests that human S1 is tuned to individual vibration frequencies (Wang and Yau, 2021). Our neural network modelling assumes that there are also neurons tuned to the amplitude of specific frequencies, though direct empirical evidence for this tuning is currently lacking. The existence of this coding would be consistent with the finding that S1 performs the initial stages of localization on a rod (Miller et al., 2019). Furthermore, resonant amplitudes are crucial pieces of information in the natural statistics of vibrations, making it plausible that they are encoded at some stage of processing. Our results therefore open up a new avenues for neurophysiological investigations into how naturalistic vibrations are encoded by the somatosensory system.

The present study demonstrates the biological possibility that the resonant feature space can stand in for the physical space of the tool, allowing for trilateration to be performed to localize touch in tool-centered coordinates. It is interesting to note that the present neural network had a similar structure to one we previously demonstrated could perform trilateration on the body surface. The biggest difference is the input layer, which must first encode the vibration information. However, once this is transformed into the representation of the feature space, the computation proceeds as it would for the body. Note that this does not necessitate that the same neural populations localize touch on limbs and tools (Schone et al., 2021), but only that the same computation is performed when localizing touch on both surfaces. Our network therefore provides a concrete demonstration of what it means to repurpose a body-based computation to localize touch on a tool. The repurposing of the neural network architecture for trilateration explains tool embodiment and the emergence of a shared spatial code between tools and skin.

## Author Contributions

L.E.M., C.F., and A.F. conceived of the behavioral experiments; C.F. performed the behavioral experiments; L.E.M. and C.F. analyzed the behavioral data; L.E.M., A.F., and W.P.M. conceived of the computational model; L.E.M and W.P.M. conceived the neural network model; F.V. and A.R. provided conceptual input; L.E.M., A.F. and W.P.M. wrote the original draft of the manuscript; All authors provided feedback on the manuscript and approved its final form.

